# Association of handgrip strength asymmetry and weakness with successful aging among older adults in China

**DOI:** 10.1101/2025.07.16.665060

**Authors:** Wei Ji, Yanping Wang, Chunping Ni, Xueyan Huang, Wenjuan He

**Author notes:** **Correspondence:** Wenjuan He, Chief Nurse, Correspondence Address:Xi’an, 710000, Shaanxi province, China. **Collaborative effort:** Both of the two authors have made equally significant contributions to the work and share equal responsibility and accountability for it (Wei Ji, Yanping Wang).

## Abstract

**Background:** Successful aging (SA) is important for the increasing population aging. The role of handgrip strength(HGS) asymmetry and weakness in successful aging remains unclear. This study aimed to elucidate the association of HGS asymmetry and weakness with successful aging in older adults.

**Methods:** We included participants aged ≥ 60 years from the 2015 China Health and Retirement Longitudinal Study (CHARLS).SA absence of major diseases, absence of major chronic diseases, no impairment in physical function, high cognitive functioning, good mental health, and active participation in life. HGS asymmetry and weakness were measured using the maximum value of the HGS. Logistic regression modeling was used to examine the association of individuals with HGS asymmetry and weakness with SA. Restricted cubic spline (RCS) modeling was used to explore potential nonlinear relationships.

**Results:** Of the 5,031 individuals included, the median age of the study population was 67 years IQR: 63-73 years, 45.6% female. Only 6.3% met the criteria for successful aging. HGS asymmetry (OR = 0.597,95 % CI: 0.472-0.754) and weakness (OR = 0.643,95 % CI: 0.417-0.964) were both independent influences on SA. Participants were less likely to have SA when both HGD asymmetry and frailty were present (OR = 0.426,95 % CI: 0.240-0.757). Further subgroup analyses revealed significant associations between HGS status and each of the components of SA, particularly with regard to physical functioning. There was an n-shaped relationship between HGS asymmetry and SA.

**Conclusion:** HGS asymmetry is associated with a reduced likelihood of weak SA. Improving or maintaining HGS symmetry and frailty may contribute to SA in older adults.

## Introduction

With the increasing trend of global population aging, how to achieve successful aging (SA) has become an important research topic in public health and social sciences [1].SA is the activities and behaviors expected of older adults who age successfully despite the inevitable losses and gradual deterioration of health that occur during aging [2]. A recent systematic evaluation and meta-analysis noted an overall estimated SA rate of 22% for people ≥60 years of age globally [3].SA not only focuses on the maintenance of physical health during the aging process but also emphasizes the enhancement of the quality of life of older adults and the preservation of psychological and social functioning during the aging process [4]. Traditionally, studies of SA have focused on the maintenance of health status, especially the maintenance of disease-free status and physical and mental functioning [2]. However, in recent years, an increasing number of studies have begun to focus on other non-biomedical factors in the aging process, such as psychological resources [4], cognitive-emotional [5], and social engagement [6], which are equally recognized as important components of successful aging.

In assessing successful aging, physical activity level, a key domain [3], is widely recognized as a composite measure of health and quality of life for older adults. A meta-analysis was conducted on the 6 aspects included more commonly in SA criteria, and the results showed that the rate of no disability was the highest [3]. Physical activity level is not only related to energy level and metabolism [7], but also closely related to muscular endurance [8], nutritional status [9], and other factors. Handgrip Strength (HGS) has been widely used in aging research as an important tool to assess muscular endurance and nutritional status [10]. Studies have shown that low grip strength values are significantly correlated with frailty [11], dysfunction [12], malnutrition [13], and the development of several diseases [14], suggesting the importance of grip strength in successful aging.

However, existing studies have focused mainly on the absolute value of grip strength (i.e., the maximum force of the grip), and less attention has been paid to the emerging metric of grip asymmetry (the significant difference in grip strength between the dominant and non-dominant hand). Grip strength asymmetry, as an important manifestation of muscle function and physical health, may reflect functional changes in the neuromuscular system during aging and may be associated with aging-related problems such as cognitive decline [15] and dysfunction [16].

In this study, nationally representative data from the China Health and Retirement Longitudinal Study (CHARLS) were used to examine the independent and pooled associations of HGS asymmetry and weakness with SA in Chinese adults ≥60 years of age. In addition, associations between HGS status and each domain of SA were investigated to identify potential interventions.

## Methods

### Study population

The source of data for this paper is the China Health and Retirement Longitudinal Study (CHARLS) Wave 3 of 2015, which provides the most comprehensive information on successful aging in recent years. CHARLS Wave3 was approved by the Biomedical Ethics Committee of Peking University (IRB00001052-11015) and the survey was conducted with the informed consent of all participants.

The CHARLS study utilizes a multi-stage sampling approach at the county, village, household, and individual levels, with probabilities proportional to size (PPS) at each stage to ensure unbiased and representative samples. CHARLS 2015 Wave 3 included 21,095 participants, and data were screened after identifying study variables, with exclusion criteria of (1) incomplete data related to the primary study variable SA, (2) less than 60 years of age, and (3) missing data related to grip strength. A total of 16,064 participants were excluded with missing data in general information and study variables. The included data were collated and analyzed, and a total of 5,031 participants were finally included(Fig. 1).

**Fig. 1.**
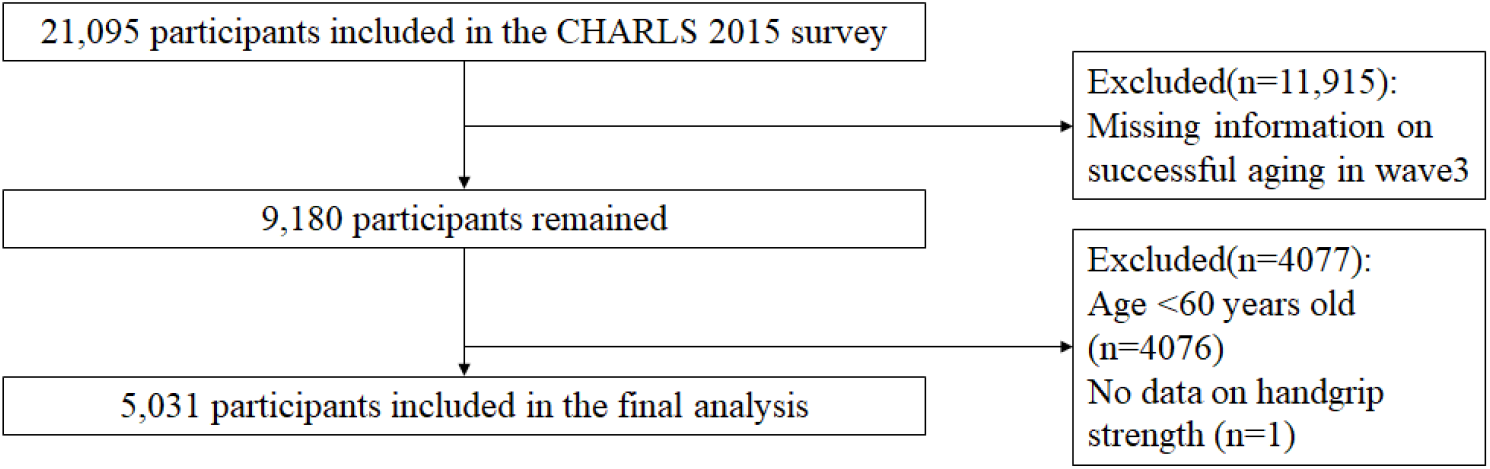
Flowchart of the study. CHARLS: China Health and Retirement Longitudinal Study; HGS: handgrip strength. wave 3:the CHARLS 2015 survey.

### Successful aging

According to Rowe and Kahn’s definition of successful aging[2], the criteria for successful aging include the following five dimensions: absence of major chronic diseases, no impairment in physical function, high cognitive functioning, good mental health, and active participation in life. Thus, participants who met all of these conditions, including the five components listed above, were categorized as being in the Successful Aging group. The single indicator of SA was operationalized as follows:

Absence of major chronic diseases: using the following series of questions: “Have you been diagnosed by a physician with any of the following medical conditions? These conditions include cancer, chronic lung disease, diabetes, heart disease, and stroke. Research has shown that the above diseases are responsible for a major burden of disease among older adults, and respondents were categorized as having no major disease if they reported no five chronic conditions[18].

No impairment in physical function: physical function was assessed using physiologically-based activities of daily living (ADL) scale. Respondents were classified as having no disability if they did not report difficulty performing any of the following six ADLs: bathing, dressing, eating, indoor transfers, toileting, or controlling urination and defecation.

High Cognitive Functioning: Participants are considered to have high cognitive functioning if they achieve median or higher scores using the Telephone Interview for Cognitive Status (TICS-10), Word Recall, and Picture Drawing. The TICS-10 consists of subtracting sequences of 7 out of 100 (up to 5 times) and correctly naming the day of the week, the month, the year, and the seasons. Word recall includes immediate and delayed recall of 10 words in a list.

Good mental health: depressive symptoms are assessed using the ten items of the Center for Epidemiologic Studies Depression Scale (CESD-10). A threshold score of ≥10 was used to identify respondents with significant depressive symptoms.

Active social participation in life: respondents were defined as active if they participated in any of the following types of social activities: volunteer or charitable work, providing assistance to family, friends, or neighbors, or participating in sports, social, or other types of clubs in the month before the interview.

### Handgrip Strength Asymmetry and Weakness

Handgrip strength (HGS) was measured using a Yuejian™ WL-1000 mechanical dynamometer (Nantong, China) as described by Zhao et al[17]. Each participant’s dominant hand was identified, and HGS was measured twice for each hand. Weakness was defined as a maximal HGS value of <28 kg for men and <18 kg for women in the dominant hand[19].

To assess HGS asymmetry, the HGS ratio was calculated by dividing the maximal HGS of the non-dominant hand by that of the dominant hand. Following the “10% rule” proposed by Armstrong and Oldham[20], which suggests that HGS in the dominant hand is typically 10% stronger than in the non-dominant hand, asymmetry was defined as an HGS ratio either <0.9 or >1.1.

Participants were then classified based on the severity of their HGS asymmetry into the following categories: Normal (ratio 0.9–1.1), Mild asymmetry (ratio 0.8–0.9 or 1.1–1.2), Moderate asymmetry (ratio 0.7–0.8 or 1.2–1.3), and Severe asymmetry (ratio <0.7 or >1.3)[21]. To evaluate the combined effects of HGS asymmetry and weakness on SA, we refer to the grouping method of the study by Li et al[21]. participants were further divided into four groups: Normal (no HGS asymmetry or HGS weakness), Asymmetry only, Weakness only, and Both (HGS asymmetry and HGS weakness).

### Covariates

The following variables were included due to their potentially confounding effects on SA: age (continuous), sex (male/female), marital status (married/unmarried), place of residence (rural/urban), and educational level (Illiterate/primary/middle school/high school+).

## Statistical analyses

Means and SDs were used for continuous variables with normal distributions; medians and interquartile ranges (IQRs) were used for non-normal distributions. Frequencies and percentages were used for categorical variables. The rank-sum test and Pearson’s chi-square test were used to compare sample characteristics between groups. Binary logistic regression models were used to examine the independent and combined associations of HGS asymmetry and weakness with both composite measures and different domains of SA. Two models were employed for each analysis: Model 1 was a crude model without adjustment; and Model 2 was adjusted for age, gender, education level, marital status, and residence. Restricted cubic spline (RCS) regression models were employed to investigate the relationship between HGS asymmetry and the Possibilities for SA.

In addition, we performed subgroup analyses to assess the robustness of the results. Given that the association between HGS and health outcomes such as multimorbidity varies by gender, we conducted a gender-stratified analysis to explore the potential heterogeneous effects of gender. All statistical analyses were performed using SPSS (version 25.0), except for the RCS regression model and forest plots, which were produced with R 4.4.3. Hypothesis testing was two-tailed, and statistical significance was set at *P* < 0.05.

## Results

### Characteristics of participants

Table 1 summarizes the characteristics of the study cohort. Among the 5,031 participants included in the study, the median age was 67 years (interquartile range [IQR]: 63-73 years), with 45.6% (n = 2,295) of participants being female. A total of 317 participants (6.3%) met the criteria for successful aging. Regarding specific indicators, 50.9% (n = 2,562) reported no major chronic diseases, 67.9% (n = 3,416) reported no physical dysfunction, 43.6% (n = 2,195) exhibited high cognitive functioning, 42.0% (n = 2,113) demonstrated good mental health, and 50.6% (n = 2,544) were actively engaged in life participation. In terms of handgrip strength (HGS) asymmetry, 2,632 individuals (52.3%) exhibited asymmetry, while 651 participants (12.9%) demonstrated weakness. The distribution of participants with mild, moderate, and severe HGS asymmetry was as follows: 1,373 (27.3%), 561 (11.2%), and 698 (13.9%), respectively. Additionally, the distribution of participants based on the presence of HGS asymmetry only, weakness only, and both asymmetry and weakness was as follows: 2,256 (44.8%), 275 (5.5%), and 376 (7.5%), respectively. Compared to individuals classified as unsuccessfully aging, those with successful aging (SA) were younger, had a higher proportion of females, were more highly educated, were less likely to be without a spouse, were less likely to reside in rural areas, and a lower prevalence of HGS asymmetry and HGS weakness (all P < 0.05, Table 1).

**Table 1.**
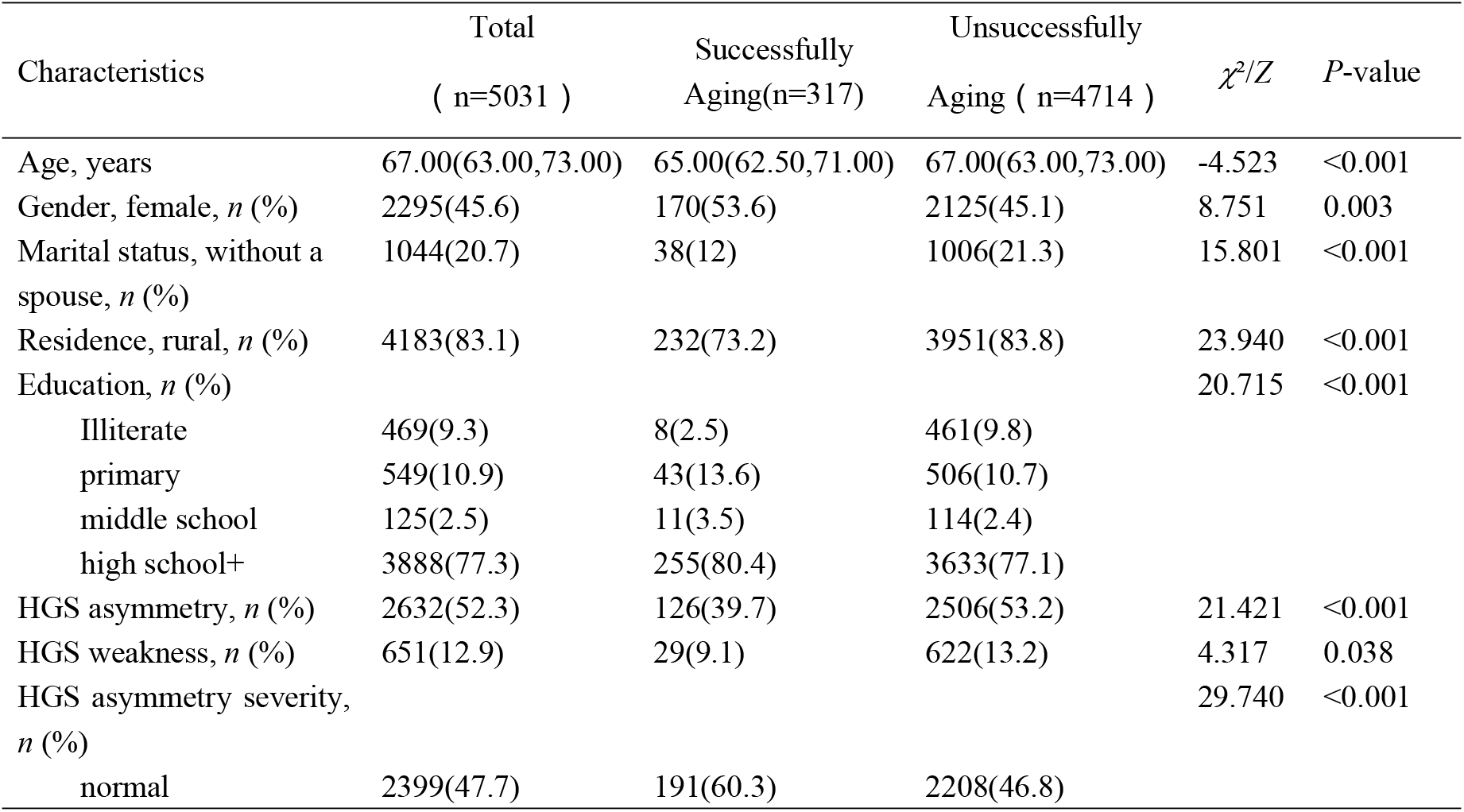
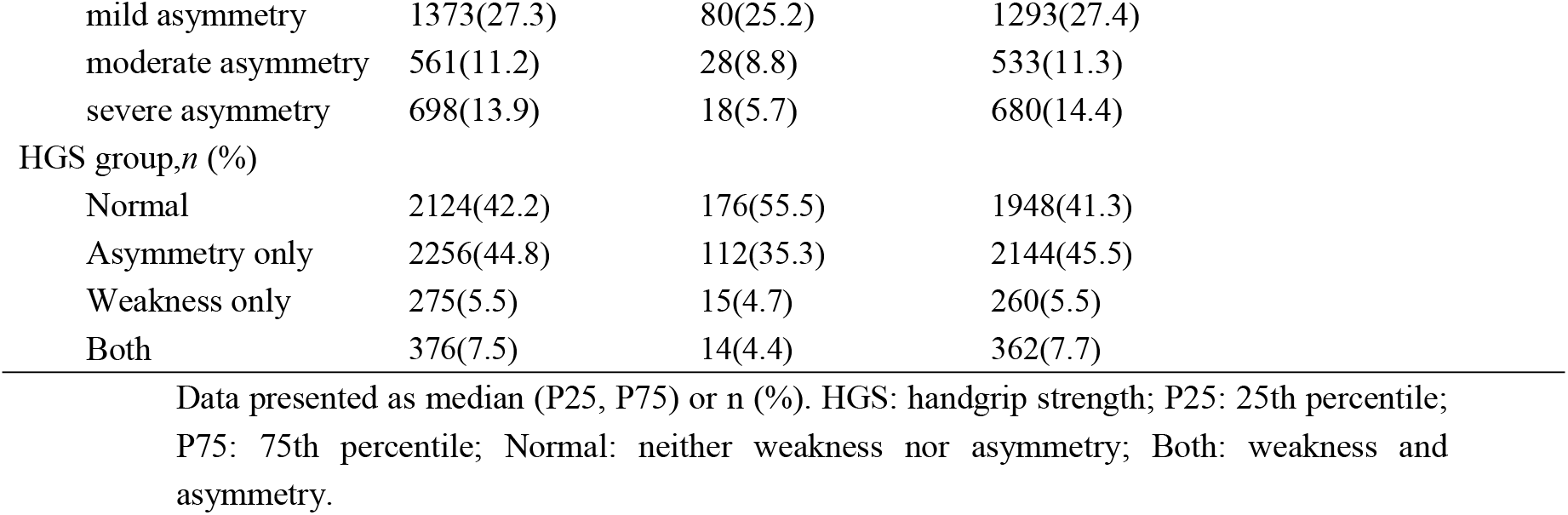
The characteristics of the participants.

| Characteristics | Total<br>( $n=5031$ ) | Successfully<br>Aging($n=317$) | Unsuccessfully<br>Aging ( $n=4714$ ) | $\chi^2/Z$ | $P$-value |
| --- | --- | --- | --- | --- | --- |
| Age, years | 67.00(63.00,73.00) | 65.00(62.50,71.00) | 67.00(63.00,73.00) | -4.523 | <0.001 |
| Gender, female, $n$ (%) | 2295(45.6) | 170(53.6) | 2125(45.1) | 8.751 | 0.003 |
| Marital status, without a spouse, $n$ (%) | 1044(20.7) | 38(12) | 1006(21.3) | 15.801 | <0.001 |
| Residence, rural, $n$ (%) | 4183(83.1) | 232(73.2) | 3951(83.8) | 23.940 | <0.001 |
| Education, $n$ (%) | | | | 20.715 | <0.001 |
| Illiterate | 469(9.3) | 8(2.5) | 461(9.8) |  |  |
| primary | 549(10.9) | 43(13.6) | 506(10.7) |  |  |
| middle school | 125(2.5) | 11(3.5) | 114(2.4) |  |  |
| high school+ | 3888(77.3) | 255(80.4) | 3633(77.1) |  |  |
| HGS asymmetry, $n$ (%) | 2632(52.3) | 126(39.7) | 2506(53.2) | 21.421 | <0.001 |
| HGS weakness, $n$ (%) | 651(12.9) | 29(9.1) | 622(13.2) | 4.317 | 0.038 |
| HGS asymmetry severity, $n$ (%) | | | | 29.740 | <0.001 |
| normal | 2399(47.7) | 191(60.3) | 2208(46.8) |  |  |

|  |  |  |  |
| --- | --- | --- | --- |
| mild asymmetry | 1373(27.3) | 80(25.2) | 1293(27.4) |
| moderate asymmetry | 561(11.2) | 28(8.8) | 533(11.3) |
| severe asymmetry | 698(13.9) | 18(5.7) | 680(14.4) |
| HGS group, *n* (%) |  |  |  |
| Normal | 2124(42.2) | 176(55.5) | 1948(41.3) |
| Asymmetry only | 2256(44.8) | 112(35.3) | 2144(45.5) |
| Weakness only | 275(5.5) | 15(4.7) | 260(5.5) |
| Both | 376(7.5) | 14(4.4) | 362(7.7) |

Data presented as median (P25, P75) or *n* (%). HGS: handgrip strength; P25: 25th percentile; P75: 75th percentile; Normal: neither weakness nor asymmetry; Both: weakness and asymmetry.

The basic characteristics of the excluded patients are as follows. In terms of age, 8,640 patients are under 60 years old, while 7,157 are 60 years old or above. The age distribution is relatively balanced. Regarding gender, there are 7,609 male patients and 8,433 female patients, with a slightly higher number of females. In terms of educational attainment, there are 1,276 illiterate patients, 1,780 with a primary education, 790 with a middle - school education, and 12,183 with a high - school education or above. The group with a high - school education or above accounts for the largest proportion. Concerning marital status, 2,204 patients are single and 13,860 are married. The married population is in the overwhelming majority. Regarding the place of residence, 12,958 patients live in rural areas and 3,106 live in urban areas. The number of patients living in rural areas far exceeds that in urban areas.

### Associations of HGS status with SA

The presence of HGS asymmetry or weakness was significantly associated with a reduced incidence of SA with OR = 0.597 (95% *CI*: 0.472-0.754), and 0.634 (95% *CI*: 0.417-0.964), respectively. Individuals with both HGS asymmetry and weakness had the lowest incidence of SA compared to the normal group OR = 0.426 (95% *CI*: 0.240-0.757). Compared to the normal group, the incidence of SA decreased as the degree of grip asymmetry became more severe, with the severe asymmetry group being the least likely to have SA in this study, OR = 0.343 (95% *CI*: 0.209-0.563).

In addition, the associations between the detailed components of HGS status and SA are shown in Figure 2.The presence of HGS asymmetry or weakness was significantly associated with a reduced likelihood of being no impairment in the physical functioning domain compared to the other four domains, with ORs of 0.599 (95% *CI*: 0.472-0.754), and 0.634 (95% *CI*: 0.417-0.964), respectively. The group with both HGS asymmetry and weakness demonstrated the same effect, with OR = 0.448 (95% *CI*:0.351,0.572).

**Table 2.** Association between HGS status and Successfully Aging.

|  | Model1 |  | Model2 |  |
| --- | --- | --- | --- | --- |
|  | OR(95%CI) | *P*-value | OR(95%CI) | *P*-value |
| HGS asymmetry(Reference = No) |  |  |  |  |
| Yes | 0.581(0.461,0.733) | **<0.001** | 0.597(0.472,0.754) | **<0.001** |
| HGS weakness(Reference = No) |  |  |  |  |
| Yes | 0.662(0.448,0.979) | **0.039** | 0.634(0.417,0.964) | **0.033** |
| HGS asymmetry severity<br>(Reference = Normal) |  |  |  |  |
| mild asymmetry | 0.715(0.546,0.937) | **0.015** | 0.722(0.550,0.948) | **0.019** |
| moderate asymmetry | 0.607(0.404,0.913) | **0.017** | 0.656(0.435,0.989) | **0.044** |
| severe asymmetry | 0.306(0.187,0.500) | **<0.001** | 0.343(0.209,0.563) | **<0.001** |
| HGS group(Reference = Normal) |  |  |  |  |
| Asymmetry only | 0.578(0.453,0.739) | **<0.001** | 0.620(0.484,0.795) | **<0.001** |
| Weakness only | 0.639(0.371,1.099) | 0.105 | 0.657(0.373,1.159) | 0.147 |
| Both | 0.428(0.246,0.746) | 0.003 | 0.426(0.240,0.757) | 0.004 |

HGS: handgrip strength; OR: odds ratio; CI: confidence interval. Model 1: a crude model; Model 2: adjusted for age, gender, education, marital status, and residence. Both: weakness and asymmetry.

**Fig. 2.**
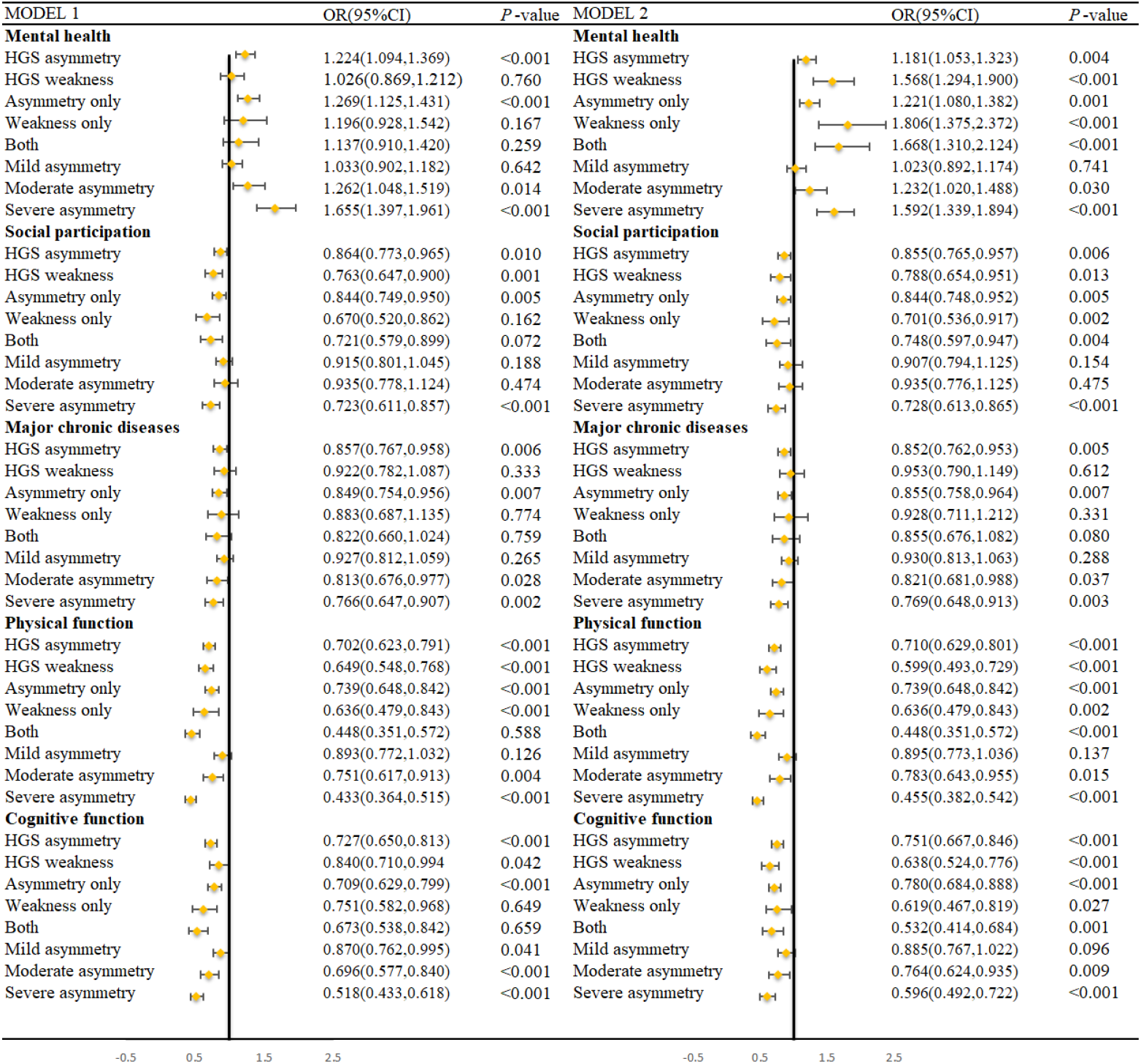
Association between HGS status and SA domains. HGS: handgrip strength; OR: odds ratio; CI: confidence interval. Both: weakness and asymmetry. Model 1: a crude model; Model 2: adjusted for age, gender, education, marital status, and residence.

Restricted cubic spline (RCS) regression with multivariable adjustment was applied to examine the dose-response relationship between the HGS ratio and the likelihood of SA, with the shaded area representing the *95%CI* (Fig. 3). As depicted in Fig. 3, a significant non-linear, n-shaped relationship was observed, indicating that a higher degree of HGS asymmetry was associated with a decreased likelihood of SA occurrence. Specifically, compared with participants without asymmetry, the odds of SA decreased with increasing severity (*OR*_mild_=0.722, 95% *CI*: 0.550–0.948; *OR*_moderate_=0.656, 95% *CI*: 0.435–0.989; and *OR*_severe_=0.343, 95% *CI*: 0.209–0.563, respectively). The two red lines indicate HGS ratios = 0.950 and 1.083, respectively, and the HGS ratio between the red lines represents a successful aging OR greater than 1.Values within this range are more strongly associated with an increased likelihood of SA.

**Fig. 3.**
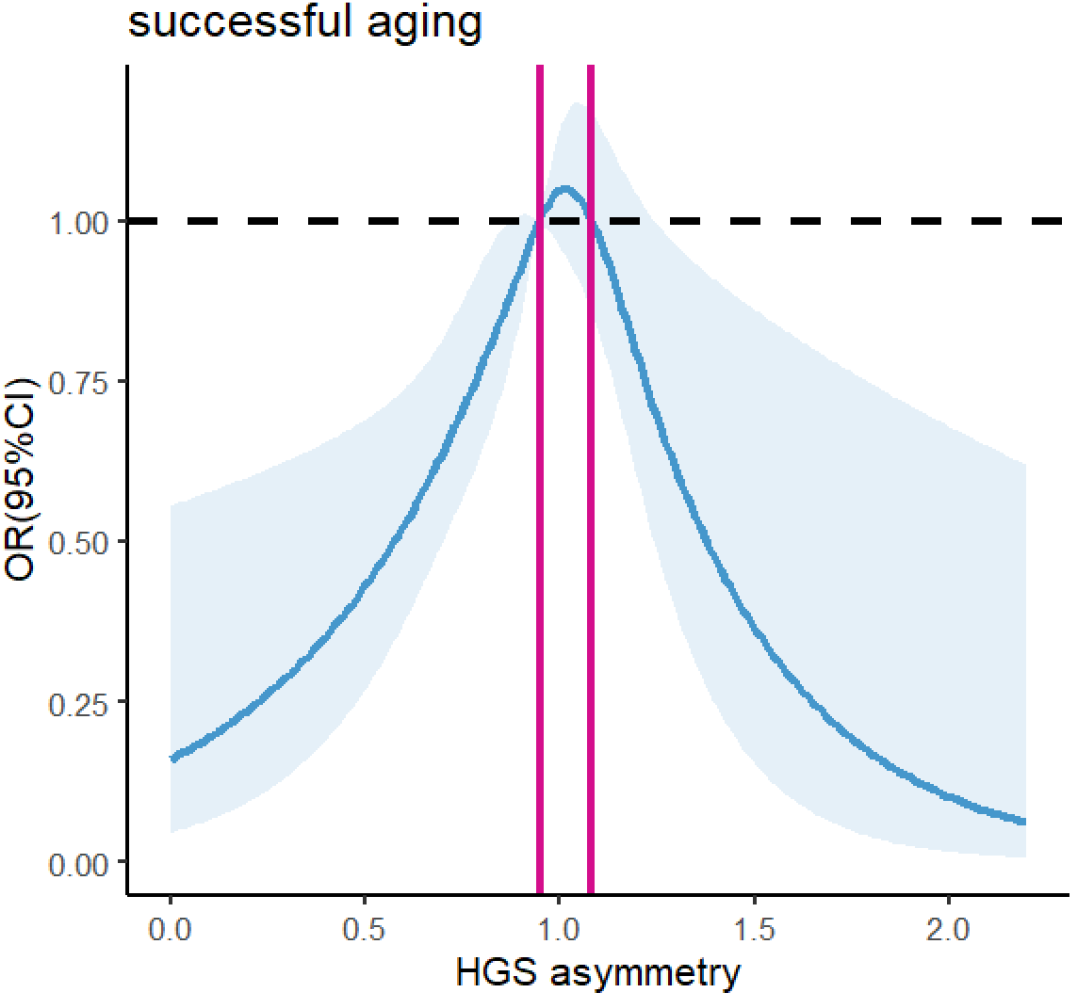
Dose-response curve of HGS asymmetry and successful aging using an RCS regression model. HGS: handgrip strength; RCS: restricted cubic spline. The two red lines indicate HGS ratios = 0.950 and 1.083, respectively, and the HGS ratio between the red lines represents a successful aging OR greater than 1. The above dose-response relationship was adjusted for following covariates: age, gender, education, marital status, and residence.

### Subgroup and sensitivity analysis

The results of the subgroup and sensitivity analyses were consistent with our main finding that HGS asymmetry and weakness were SA influencing factors. Supplementary Table S1 presents the results stratified by gender. For men, after adjusting for all covariates, except for ‘Weakness only’, ‘Both’, and ‘Moderate asymmetry’, the rest of the results were significant with SA, and for the female group only the ‘HGS asymmetry’, ‘Asymmetry only’ and ‘Severe asymmetry’ results were significant.

## Discussion

In this study, we found that both HGS asymmetry and frailty were significantly associated with a reduced likelihood of SA in Chinese older adults, with the lowest likelihood of SA in the presence of both conditions. Furthermore, we observed significant associations between HGS status and each domain of SA. These findings emphasize the importance of assessing HGS asymmetry and weakness to identify older adults with a lower likelihood of SA and associated adverse health outcomes.

Consistent with previous studies, handgrip strength asymmetry or weakness is strongly associated with both impaired physical and cognitive functions [12,16,22,23]. Furthermore, at the biological level, mitochondrial dysfunction in muscle cells, reduced muscle protein synthesis, and declines in key hormones such as testosterone and growth hormone are significant factors influencing grip strength in older adults [22,24]. As muscle mass decreases, there is a concomitant reduction in grip strength, which primarily impacts the physical mobility of older individuals[18]. Therefore, preventive and management strategies focusing on the preservation of grip strength and addressing imbalances could offer multiple benefits. Such interventions not only improve energy metabolism but also enhance vitality and have the potential to positively affect other aspects of sarcopenia (SA). This is particularly critical for the elderly population suffering from significant grip imbalances, as this study observed the lowest likelihood of sarcopenia in individuals with normal grip strength compared to those with impaired grip strength.

In this study, we found that HGS asymmetry and weakness were strongly associated with SA in Chinese older adults. Additionally, HGS asymmetry was linked to all domains of SA. First, our findings align with previous studies reporting that declines in HGS predict physical disability and loss of independence [16]. HGS asymmetry has been associated with motor dysfunction, particularly slower gait speed and poorer postural balance [21]. Furthermore, HGS asymmetry may reflect imbalanced hemispheric activation or impaired neurological function [25]. Numerous studies have shown that HGS asymmetry increases the risk of cognitive decline [19,26]. Second, HGS asymmetry is strongly associated with mental health, particularly depression. Previous studies have demonstrated that both HGS asymmetry and low HGS are linked to depression, with a stronger association when both factors are present [21,27]. This may be due to the impact of asymmetrical hemispheric activation on cognitive processes such as language, spatial attention, and social perception [28]. Moreover, HGS asymmetry may affect social participation levels. This association could be explained by the relationship between HGS asymmetry and diminished physical functioning [12], as reduced physical ability may lead individuals to decrease their social engagement. Finally, HGS asymmetry may influence the occurrence of major diseases in older adults. Consistent with previous research [29], both HGS weakness and asymmetry may increase the risk of cardiovascular outcomes in elderly individuals. Since this study did not investigate specific diseases, further research is needed to explore the potential relationship between grip asymmetry and particular major diseases in older adults.

Finally, we also found that the likelihood of successful aging was lowest in individuals with both grip asymmetry and weakness, with particularly pronounced results in males when performing gender subgroup analyses. The prevalence of handgrip strength (HGS) asymmetry was more than twice as high as that of weakness, suggesting that HGS asymmetry may represent an early form of muscle dysfunction that precedes the decline in overall grip strength [30]. An HGS ratio between 0.950 and 1.083 was associated with a higher likelihood of successful aging (odds ratio > 1), further refining the range of values for HGS asymmetry that is relevant when analyzing SA. However, several limitations must be considered. First, interpreting causal relationships between HGS state and SA is challenging due to the cross-sectional nature of the study design. Second, female participants with HGS weakness were not included in the final study population, which resulted in the inability to observe gender differences consistent with subgroup analysis findings. Third, as in other studies [31], diagnoses of major diseases were based on self-reported physician diagnoses. Medical records were not available in CHARLS, and therefore, the reliance on self-reported chronic disease diagnoses may introduce some degree of reporting bias. Despite these limitations, this study offers valuable insights into the underexplored relationship between grip asymmetry, weakness, and successful aging in the Chinese elderly population, contributing to a better understanding of these factors in aging research.

## Conclusions

This study suggests that the presence of HGS asymmetry and frailty is independently associated with a reduced likelihood of SA, and that HGS asymmetry is independently associated with each component of SA, with the lowest likelihood of SA occurring when both are present. Maintaining HGS symmetry and reducing asymmetry may help promote SA in older adults.

## Glossary

HGS: hand grip strength
SA: successful aging
RCS: Restricted cubic spline

## Appendices

Supplementary Table S1. Association between HGS status and SA in different gender groups.

Supplementary Fig.S1. Dose-response curve of HGS asymmetry and successful aging using an RCS regression model.

## Acknowledgements

We thank Peking University for the open data resources and all investigators who participated in the study.

## Author contributions

Wei Ji and Yanping Wang created the study protocol, performed the statistical analyses, and wrote the first manuscript draft. Chunping Ni confirmed the data and assisted with the statistical analyses. Xueyan Huang contributed to data interpretation and manuscript revision.Wenjuan He, as senior author, reviewed and edited all versions of this manuscript. All authors read and approved the final manuscript.

## Availability of data and materials

The data for this article comes from the the China Health and Retirement Longitudinal Study database for 2015. Available from https://charls.pku.edu.cn/en/.

## Declarations

### Clinical trial number

not applicable

### Ethics approval and consent to participate

This is a retrospective study based on CHARLS database. The patient’s information has been hidden before the study. The original CHARLS was approved by the Ethical Review Committee of Peking University (IRB00001052–11015), and all participants signed the informed consent at the time of participation. This research followed the guidance of the Declaration of Helsinki.

## Declaration of Generative AI and AI-assisted technologies in the writing process

During the preparation of this work, the first author used DeepL Write to improve the manuscript’s readability. After using this tool, the author reviewed and edited the content as needed and takes full responsibility for the content of the published article.

